# Human CD4^+^/CD8α^+^ regulatory T cells induced by *Faecalibacterium prausnitzii* protect against intestinal inflammation

**DOI:** 10.1101/2022.03.22.485277

**Authors:** Sothea Touch, Emmanuelle Godefroy, Nathalie Rolhion, Camille Danne, Cyriane Oeuvray, Marjolène Straube, Chloé Galbert, Loïc Brot, Iria Alonso Salgueiro, Sead Chadi, Tatiana Ledent, Jean-Marc Chatel, Philippe Langella, Francine Jotereau, Frédéric Altare, Harry Sokol

**Affiliations:** Sorbonne Université, INSERM, Centre de Recherche Saint-Antoine, CRSA, AP-HP, Saint Antoine Hospital, Gastroenterology department, F-75012 Paris, France; Paris Center for Microbiome Medicine (PaCeMM) FHU, Paris, France; CRCINA, INSERM, University of Nantes, University of Angers, Nantes, France; INRAE, UMR1319 Micalis & AgroParisTech, Jouy en Josas, France

**Keywords:** Tregs, *Faecalibacterium prausnitzii*, inflammatory bowel diseases, microbiota, inflammation, DSS-induced colitis, DP8α

## Abstract

*Faecalibacterium prausnitzii (F. prausnitzii)*, a dominant bacterium of the human microbiota, is decreased in patients with inflammatory bowel diseases (IBD) and exhibits anti-inflammatory effects. In human, colonic lamina propria contains IL-10-positive, Foxp3-negative regulatory T cells (Treg) characterized by a double expression of CD4 and CD8α (DP8α) and a specificity for *F. prausnitzii*. This Treg subset is decreased in IBD. The *in vivo* effect of DP8α cells has not been evaluated yet. Here, using a humanized model of NOD.*Prkcscid IL2rγ^-/-^* (NSG) immunodeficient mouse strain that expresses human leucocyte antigen D-related 4 (HLA-DR4) but not murine class II (NSG-Ab° DR4), we demonstrated a protective effect of DP8α Tregs combined with *F. prausnitzii* administration in a colitis model. In a cohort of patients with IBD, we showed an independent association between the frequency of circulating DP8α cells and disease activity. Finally, we pointed out a positive correlation between *F. prausnitzii*-specific DP8α Tregs and the amount of *F. prausnitzii* in fecal microbiota in healthy individuals and patients with ileal Crohn’s disease.

## INTRODUCTION

Inflammatory bowel diseases (IBD) include Crohn’s disease (CD) and ulcerative colitis (UC) and are characterized by chronic and relapsing inflammation of the intestine. Their incidence rates and prevalence are high and rising in Western countries. Identifying markers to predict relapse or complications and new treatments are much needed and would have high socio-economic impacts. The exact pathogenesis of IBD remains to be deciphered. However, it involves dysregulated immune responses to commensal bacteria (1) and the implication of genetic factors as emphasized by genome-wide association studies (GWAS) and mouse models (2–5). In addition, environmental factors and life habits are also involved and many studies have highlighted the role of gut microbiota alterations in the pathogenesis (6, 7). One of the strongest signal repeatedly identified in the IBD-associated microbiota alteration is the decreased abundance of *Faecalibacterium prausnitzii* (*F. prausnitzii*), a dominant bacterium of the *Clostridium* IV group (8–10). *F. prausnitzii* is highly prevalent in the normal human gut microbiota and represents by itself around 5% of the total bacterial population in healthy individuals (11). We and others showed that it exerts anti-inflammatory properties both *in vitro* and *in vivo* in different colitis mouse models (12–14). More than a simple witness of inflammation, the role of the gut microbiota as an actor in IBD pathogenesis is now demonstrated. Among the strongest arguments, one can cite the pro-inflammatory effect of the IBD-associated microbiota demonstrated by microbiota transfer from human to mice (15), and the clinical efficacy of fecal microbiota transplantation as a treatment of IBD (16). In mice, *Clostridium* IVa and XIV bacteria play a role in inducing interleukin (IL)-10-secreting forkhead box P3 (Foxp3)-positive regulatory T cells (Tregs) in the colonic lamina propria (LP), which attenuate colitis (17). In humans, the proportion of Foxp3^+^ Tregs is low in the colonic LP, as compared to mice (5% and 25% respectively) (18, 19) and despite the extensive interest in Foxp3^+^ Tregs as key players in the intestinal immune homeostasis, there is little evidence for the role of these Tregs in IBD. No defect in the abundance or function of these cells was observed in IBD patients (19, 20) and no polymorphisms in Foxp3 or other genes related to Foxp3^+^ Treg differentiation was detected by GWAS. In addition, individuals with immune dysregulation, polyendocrinopathy, enteropathy, X-linked (IPEX) syndrome and thus lacking functional Foxp3^+^ Tregs, do not necessarily develop colitis (21), supporting the hypothesis that Foxp3^+^ Tregs are not essential in the development of IBD. In this context, we have recently characterized, in the human colonic LP, a subset of Foxp3-negative IL-10-producing Tregs responding to *F. prausnitzii* in a major histocompatibility complex (MHC) class II and T cell receptor (TCR)-dependent manner (22, 23). These cells share most *in vitro* regulatory functions with Foxp3^+^ Tregs and co-express CD4 and low levels of CD8α *in vivo,* hence were named double-positive CD8α (DP8α). We found that DP8α Tregs are abundant in the healthy colonic LP (up to 13% of CD4^+^ T cells). They are also present in blood, where CCR6^+^/CXCR6^+^ DP8α cells react to *F. prausnitzii,* in contrast to DP8α cells lacking one or both receptors (22), the role of which remains unknown. Interestingly, both colonic and circulating DP8α Tregs are drastically reduced in IBD patients, as compared to healthy subjects (22). Along with these findings, we have recently revealed the ability of *F. prausnitzii* to induce tolerogenic dendritic cells that favor IL-10-producing CD4 T cells priming *in vitro* (24). Altogether, these data argue that *F. prausnitzii* contributes to the induction of IL-10-producing DP8α Tregs in a way similar to the induction of IL-10^+^ Foxp3^+^ Tregs by murine *Clostridia* (25). This also suggests the existence of a link between reduced levels of *F. prausnitzii* and decreased activity of *F. prausnitzii*-specific DP8α Tregs in IBD patients, potentially resulting in disturbed gastro-intestinal homeostasis. Consequently, this suggests that DP8α Tregs could play a role in the control of IBD. Here, we aimed at assessing the role of DP8α Tregs *in vivo* with a sophisticated pre-clinical model. We used immunodeficient mice, humanized for the DRb1*0401 HLA class II allele and injected them with human HLA-DRb1*0401^+^ CD4^+^ T cells in the absence or in the presence of human HLA-DRb1*0401-restricted DP8α Tregs, before triggering dextran sulfate sodium (DSS)-induced colitis. In addition, we analyzed the phenotype and frequency of DP8α cells and the abundance of *F. prausnitzii* in the fecal microbiota in a large cohort of healthy controls and patients with IBD.

Here, we demonstrate the capacity of DP8α Tregs to attenuate colitis severity in a humanized mouse model, supporting their importance in the maintenance of gastro-intestinal homeostasis and in IBD. In patients with IBD, we show that a low abundance of blood DP8α cells is independently associated with several clinical parameters of disease activity such as flare and elevated C-reactive protein. Compared to healthy controls, we also confirm that patients with IBD exhibit a decreased abundance of CCR6^+^/CXCR6^+^ DP8α Tregs and of *F. prausnitzii* in their blood and fecal microbiota, respectively. Interestingly, both parameters were positively correlated in CD patients with ileal involvement, suggesting that *F. prausnitzii* could induce DP8α Tregs in humans, in the same manner *Clostridia* induce Foxp3-positive Tregs in mice. These data extend the pathologic relevance of the decreased abundance of DP8α cells and suggest that DP8α Treg abundance may both serve as an indicator of intestinal inflammation and represent a therapeutic target in IBD.

## RESULTS

### Validation of the mouse model and adoptive transfer of human cells

To evaluate the effect of DP8α Tregs on intestinal inflammation *in vivo,* we used a model of dextran sulfate sodium (DSS)-induced colitis in non-obese diabetic (NOD).Prkdcscid.Il2rg-/- (NSG) mice that lack murine major histocompatibility complex (MHC) class II and instead can express a human leukocyte antigen (HLA) allele of interest under the control of the murine MHC class II promoter as described before (26). NSG mice lack murine T cells, B cells and natural killer (NK) cells and can be reconstituted with human CD4^+^ T cells from allelically-matched HLA-class II donors. We used NSG mice expressing the frequently expressed DRb1*0401 allele (NSG-Ab°DR4 mice), reconstituted with purified HLA-DRb1*0401-expressing CD4^+^ T cells. Then to assess the role of DP8α Tregs in intestinal inflammation, we set up a colitis model in these mice and generated DP8α Tregs responding to HLA-DRb1*0401-expressing mouse antigen presenting cells loaded with *F. prausnitzii*.

To select HLA-DRb1*0401-restricted DP8α Tregs responding in NSG-Ab°DR4 mice, we used PBMCs from two donors of the blood bank (Etablissement Français du Sang, EFS, Pays de Loire, France), one typed homozygous for the relevant DRb1*0401 allele and the second, used as a control, was DRb1*0408/DRb1*1396. CD4^+^ T cells were magnetically-sorted and DP8α Tregs were FACS-sorted and cloned simultaneously before being expanded using polyclonal stimulation as we described before (22). CD4^+^ T cell purity was confirmed after amplification (> 98% purity). We first selected DP8α Treg clones best responders to *F. prausnitzii-loaded* autologous monocytes and then, among these, a clone from each donor whose response was specifically inhibited by a blocking anti-HLA DR antibody (Supplemental Figure 1A-B). Finally, we confirmed that murine bone marrow derived dendritic cells (BM-DCs) from NSG-Ab°DRb1*0401^+^ mice loaded with *F. prausnitzii* induced a potent cytokine response (IL-10 and IFN-ɣ) from the selected DRb1*0401 homozygous clone, but not from the DRb1*0408/DRb1*1396 one (Supplemental Figure 1C-F), validating not only the HLA-DRb1*0401-restricted response to *F. prausnitzii* of the first clone but also, importantly, its ability to respond as well to BM-DCs from NSG-Ab°DR4 mice.

We first set up the DSS-induced colitis model in a preliminary experiment (data not shown), in which mice received intraperitoneal injection of human allelically-matched effector CD4^+^ T cells ten days before DSS administration. We determined that the dose of 1% DSS in drinking water was suitable to induce weight loss and colitis without causing too much mortality and allow recovery upon DSS removal.

We used flow cytometry, to evaluate the presence of human cells in NSG-Ab°DR4 mice after intraperitoneal injection of CD4^+^ effector T cells in combination or not with allelically-matched DRb1*0401-restricted DP8α. This was evaluated in the blood ten days after T cell injection before the induction of DSS-induced colitis and in the colonic LP, 18 days after T cell injection and 8 days after the start of DSS treatment, to ensure T cell recruitment to the intestinal mucosa of mice (Figure 1). Cells were gated according to side scatter (SSC-A) and forward scatter (FSC-A) after doublet discrimination. The frequency of human cells was then measured as the fraction of human CD45^+^/mouse CD45^-^ cells among total, mouse and human, CD45^+^ cells. In the colon LP, human CD3^+^/mouse CD45^-^ cells were gated among total mouse CD45^+^ and human CD3^+^ live cells. Human cells were virtually all CD3^+^/CD4^+^ T cells both in blood (Figure 1B-C) and in the colon LP (Figure 1D-E). The frequency of human T cells in the blood and colon LP was not significantly different between CD4 + PBS and CD4 + DP8α groups (Figures 1B and 1D). We could not distinguish CD4^+^ effector T cells from DP8α clones, as most DP8α clones lose CD8α expression after *in vitro* expansion, without any impact on their *in vitro* regulatory functions (unpublished results). To address this issue and confirm the presence of DP8α clone in NSG-AB°DR4 mice, we injected a group of mice with DP8α cells only in absence of CD4^+^ cells and compared it to vehicle-treated mice. After 10 days of gavage with *F. prausnitzii,* the mice received 1% DSS in drinking water for 7 days (Figure 1F) and the frequency of human CD3^+^ cells in the colon lamina propria was quantified by flow cytometry at different timepoints (Figure 1G). We show an engraftment of human DP8α clone in the colonic mucosa that is increasing with time. Altogether, we hereby confirmed the presence of human T cells in the blood (1.5 to 5.5% of total mouse and human CD45^+^ cells) and in the colon LP of inflamed mice (0.57 to 3.8% of total mCD45^+^ and hCD3^+^ cells), thus validating our model and allowing further investigations.

**Figure 1.**
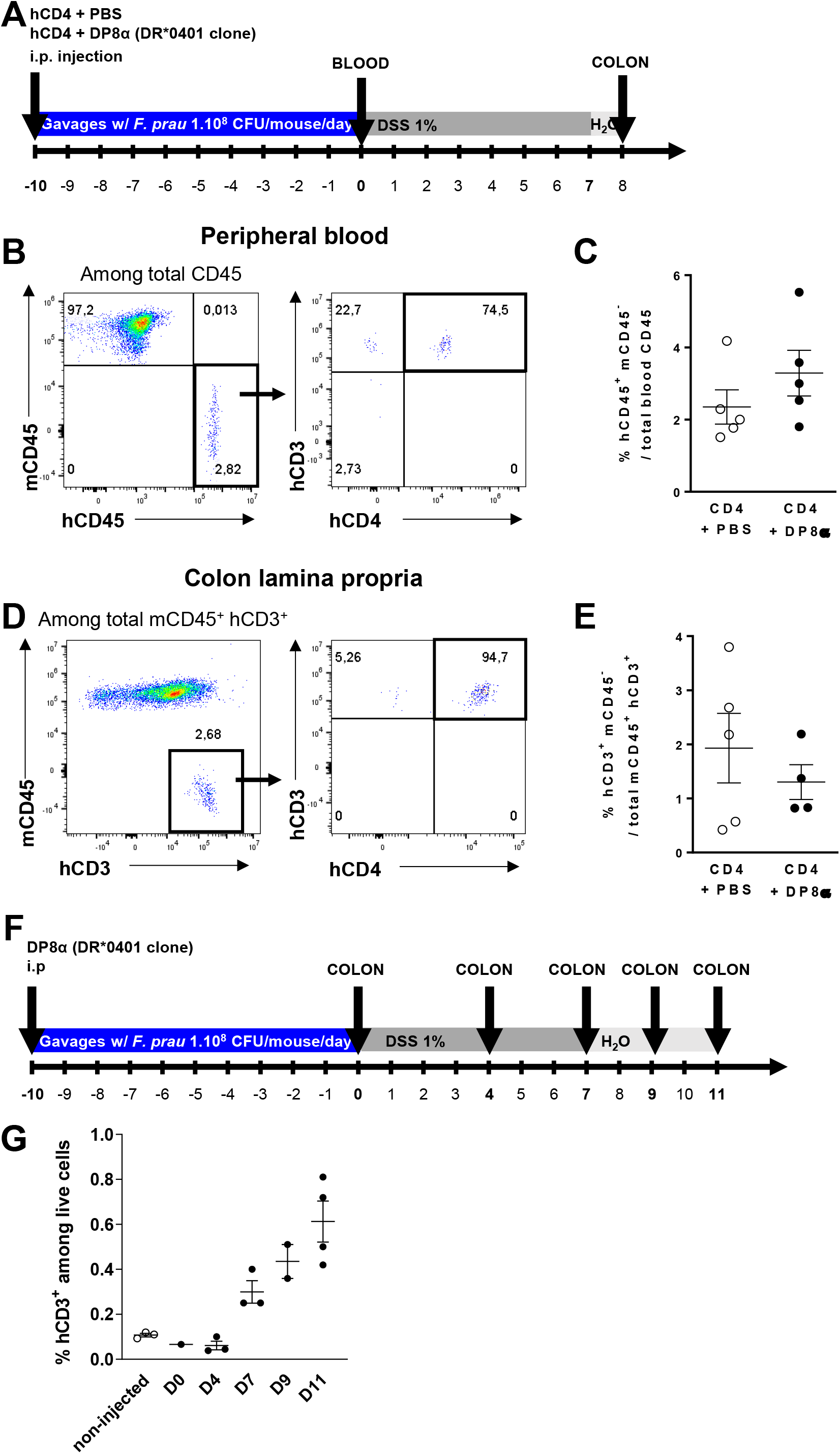
Detection of human cells in whole blood and the colon lamina propria of NSG-Ab°DR4. **(A)** The presence of human cells after injection of CD4 effector T cells in presence or absence of DP8α Tregs was assessed by flow cytometry in two independent experiments in the peripheral blood at day 0 and in the colon lamina propria at day 8 after DSS treatment in CD4 + PBS (n = 5 and n = 5), empty dots) and CD4 + DP8α (n = 4 and n = 5, black dots) groups. **(B)** After doublet discrimination and exclusion of debris, human CD3^+^ and human CD4^+^ were analyzed among human CD45^+^ murine CD45^-^ cells. **(C)** Frequency of human CD45^+^ mouse CD45^-^ cells among total CD45^+^ cells. **(D)** Detection of human cells in the colon lamina propria was determined by the expression of human CD3 and the absence of expression of mouse CD45 among total mCD45^+^ and hCD3^+^ cells. **(E)** Frequency of human CD3^+^ mouse CD45^-^ cells among total colonic mCD45^+^ and hCD3^+^ cells. **(F-G)** The presence of DP8α clone was evaluated by flow cytometry after 10 days of gavage with *F. prausnitzii* at day 0, 4, 7, 9 and 11 after the start of DSS treatment. Non-injected indicates the control group without cells injected. Results are presented as the mean ± S.E.M.

### Administration of DP8α cells with F. prausnitzii protects against DSS-induced colitis

Next, we studied the ability of human DRb1*0401-restricted DP8α Tregs, alone or in combination with *F. prausnitzii* intragastric gavage, to alleviate DSS-induced colitis in NSG-Ab°DR4 mice. The mice were injected intraperitoneally with human CD4 peripheral T cells and PBS or DP8α clone and received PBS or *F. prausnitzii* by gavage for 10 days before the start of DSS treatment for 7 days followed by a recovery phase of 4 days with regular water (Figure 2A). As shown by assessing body weight loss, disease activity index (DAI) and survival, DP8α Tregs combined with *F. prausnitzii* exhibited protective effects, while the administration of DP8α Tregs alone did not (Figure 2B-C). The histological score, evaluating the severity, spread of inflammation and crypt damage, was significantly lower in CD4 + DP8α Tregs + *F. prausnitzii* group compared to CD4 + PBS and CD4 DP8α (Figure 2D-E), confirming a protective effect of activated DP8α Tregs.

**Figure 2.**
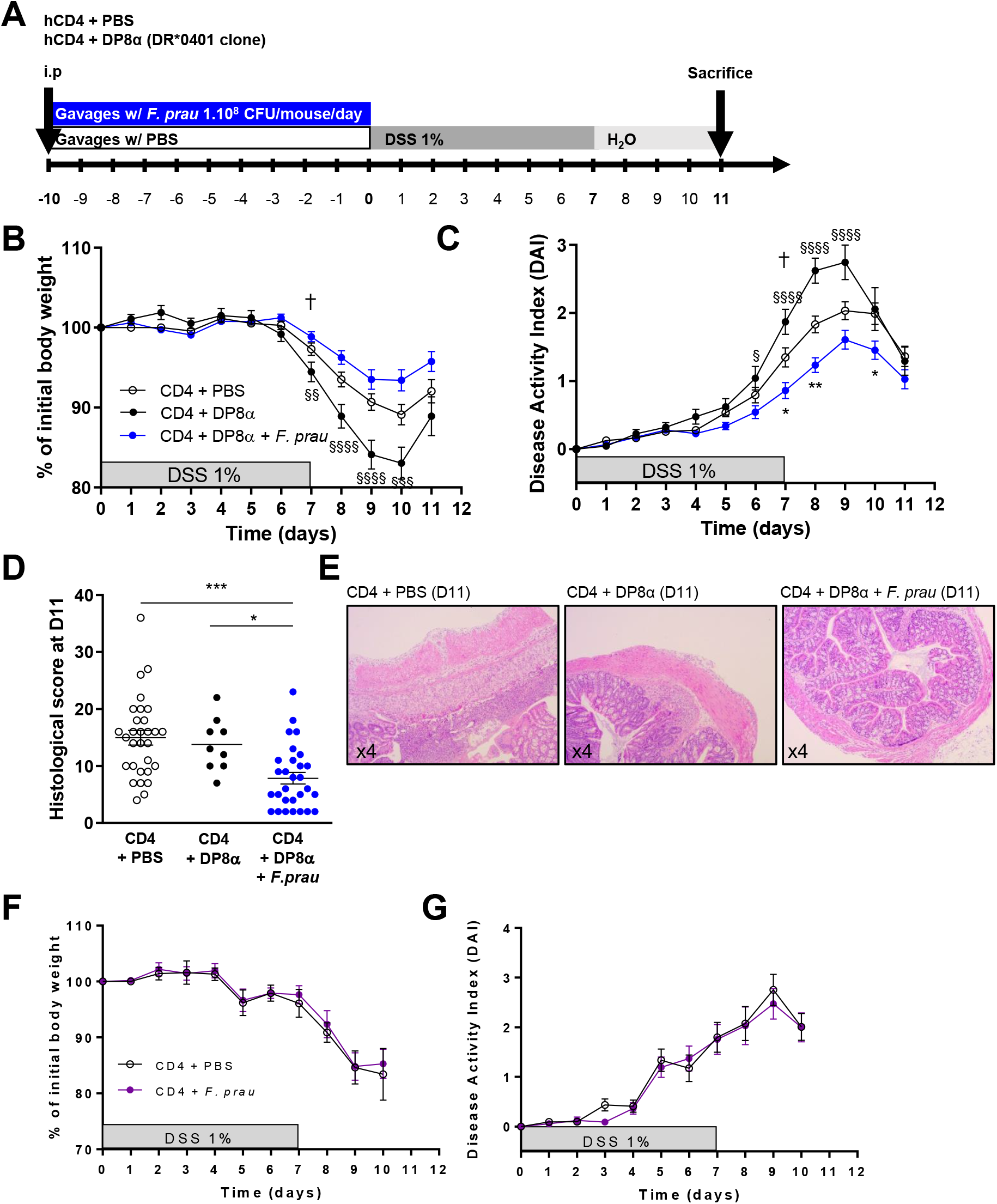
DP8α Tregs + *F. prausnitzii* administration, but not DP8α or *F. prausnitzii* alone protect NSG-Ab°DR4 mice from DSS-induced colitis. **(A)** Experimental outline: NSG-Ab°DR4 female mice were injected intraperitoneally (i.p.) with PBS or human peripheral CD4 effector T cells with either a human DRb1*0401/DRb1*0401 DP8α Treg clone or PBS and received daily gavage with 1 x10^8^ CFU of *F. prausnitzii* or 200 μl vehicle 1X PBS for 10 days before 1% DSS supplementation in drinking water for 7 days followed by 4 days of regular drinking water. **(B)** Body weight and **(C)** DAI were evaluated in CD4 + PBS (n = 44), CD4 + DP8α (n = 20) and CD4 + DP8α + *F. prausnitzii* (n = 45) groups in 5 independent experiments. (**D-E**) Histological score was assessed in CD4 + PBS (n = 30), CD4 + DP8α (n = 9) and CD4 + DP8α + *F. prausnitzii* (n = 29) groups in 3 independent experiments. **(F)** Body weight and **(G)** DAI were also evaluated in CD4 + PBS (n = 10, open circles), as compared to CD4 + *F. prausnitzii* groups (n = 11, purple circles) in another series of 2 independent experiments. Body weight and DAI are presented as the mean ± S.E.M. **P < 0.01, and § P<0.05, §§ P < 0.01 by one-way analysis of variance (ANOVA) followed by Tukey’s multiple comparison tests. The symbol † indicates the beginning of mortality.

Importantly, the intragastric administration of *F. prausnitzii* alone, without DP8α cells, did not have any protective effect, ruling out an independent effect of the bacterium (Figure 2F-G). Moreover, there was no protective effect of DP8α Tregs combined with *F. prausnitzii* when using DR-restricted DRb1*0408/DRb1*1396-expressing DP8α Tregs (Supplemental Figure 2A-C), which did not recognize *F. prausnitzii* on DRb1*0401^+^ mouse dendritic cells (Supplemental Figure 1D and 1F).

### DP8α cells administered with F. prausnitzii decrease gut inflammation and ameliorate histological score in DSS-induced colitis in mice

To further compare mice receiving CD4 + DP8α Tregs + *F. prausnitzii* versus CD4 + PBS control mice, we sacrificed mice and collected samples at D7 after DSS initiation to evaluate acute colon inflammation severity (Figure 3A).

**Figure 3.**
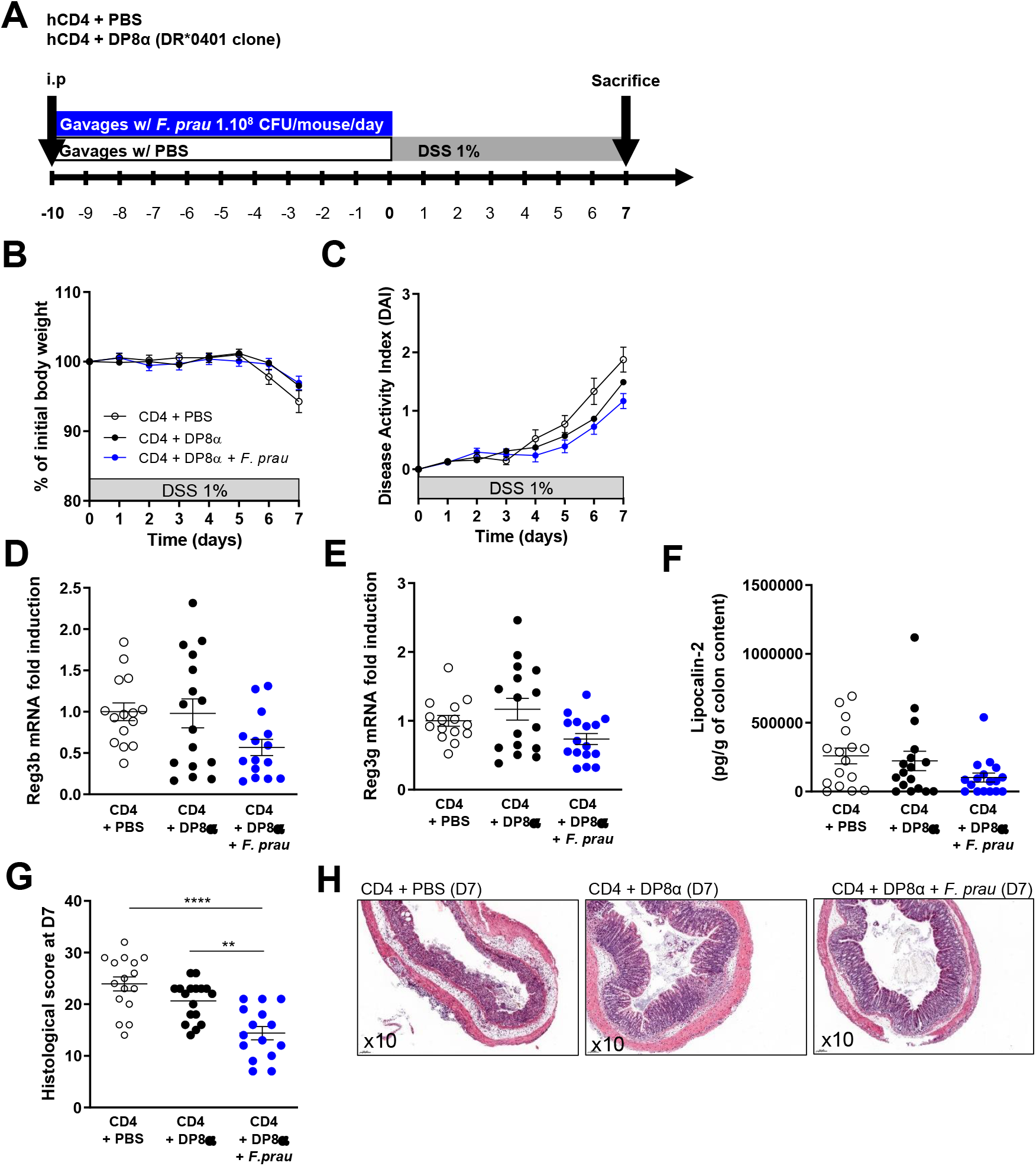
DP8α Tregs + *F. prausnitzii* administration lowers inflammation induced by DSS-induced colitis in NSG-Ab°DR4 mice. **(A)** Experimental outline: NSG-Ab°DR4 female mice were injected intraperitoneally (i.p.) with 2.10^6^ human peripheral CD4 effector T cells alone or in combination with 2.10^6^ human DRb1*0401/DRb1*0401 DP8α clones and received daily gavage with 200 μl PBS 1X or 1.10^8^ CFU of *F. prausnitzii* respectively for 10 days before 1% DSS supplementation in drinking water for 7 days. Mice were sacrificed at D7). **(B)** Body weight and **(C)** disease activity index (DAI) were assessed during the protocol in all groups of mice. **(D, E)** mRNA levels of *Reg3b* and *Reg3g* were analyzed by RT-qPCR in the proximal colon at D7, **(F)** lipocalin-2 secretion (pg/g of colon content) was measured by ELISA in the colon content at D7. Histological score was obtained from the colon of mice at D7 **(G-H)**. Results are presented as the mean ± S.E.M. For comparison between multiple groups, one-way analysis of variance (ANOVA) was performed, and P value < 0.05 were considered significant (*P <0.05, **P < 0.01, ***P < 0.001, ****P < 0.0001 and § P<0.05, §§ P < 0.01). Only significant statistical results after adjustment for False discovery rate are shown (Benjamini Hochberg, q< 0.1). Each figure is representative of N = 3 independent experiments (CD4 + PBS, n = 17; CD4 + DP8α, n = 16; CD4 + DP8α + *F. prausnitzii,* n = 17)).

At D7, no statistically significant difference was observed regarding body weight loss (Figure 3B). However, statistically significant protection was observed regarding DAI in the CD4 + DP8α Tregs + *F. prausnitzii* group (Figure 3C). Due to the important immunodeficiency of NSG mice, the numerous cytokines we explored, namely murine and human *Il-10, IL-1b, tumor necrosis factor-α, IL-17A* and *IL-22,* were either weakly expressed and not significantly different among the study groups or not detected (data not shown). Murine *Cxcl1, Cxcl2 and Cxcl5,* expressed by innate immune cells, were detected by RT-qPCR in the distal and proximal colon but no statistically significant difference was observed between CD4 + DP8α Tregs + *F. prausnitzii* and CD4 + PBS groups (data not shown). However, we showed that mRNA expression of antimicrobial C-type lectin molecules regenerating islet-derived protein 3 beta (*Reg3b*) and 3 gamma (*Reg3g*) were significantly decreased in the proximal colon of CD4 + DP8α + *F. prausnitzii* mice when compared to CD4 + PBS mice whereas no difference was observed with CD4 + DP8α group (Figure 3D-E). At the protein level, a lower abundance of the intestinal inflammation marker lipocalin-2 (Lcn-2) was observed in the total colon content of mice treated with DP8α + *F. prausnitzii*, as compared to controls (Figure 3F). The histological scorewas significantly lower in CD4 + DP8α Tregs + *F. prausnitzii* group, confirming a protective effect of activated DP8α Tregs (combination of DP8α + *F. prausnitzii)* and not DP8α alone (Figure 3G-H).

Altogether these data show that the administration of DP8α Tregs together with *F. prausnitzii* ameliorates DSS-induced colitis in humanized mice, emphasizing the role of *F. prausnitzii-* activated DP8α cells to protect against intestinal inflammation.

### The number of circulating CCR6^+^/CXCR6^+^ DP8α Tregs and the quantity of F. prausnitzii in fecal microbiota are positively correlated in healthy controls and CD patients with ileal involvement

To evaluate the translational relevance of our findings in mice, we took advantage of a large cohort of 250 patients with IBD (185 with CD and 65 with UC, clinical characteristics are described in Supplemental Table 1 and Supplemental Table 2) and 73 healthy controls (HC). Human peripheral blood mononuclear cells (PBMCs) were isolated and analyzed by flow cytometry to evaluate the frequency and phenotype of blood DP8α cells. DNA from stool samples was extracted in a subcohort of 52 IBD patients and 10 HC to quantify *F. prausnitzii* in the fecal microbiota.

Multivariate logistic regression identified several clinical parameters independently associated with the abundance of DP8α in peripheral blood (Table 1). Interestingly, low frequency (< median) of DP8α cells was associated with parameters of disease activity including flare and elevated C-reactive protein (CRP, > 5), supporting a potential anti-inflammatory role of these cells. The other factors associated with low frequency of DP8α cells included previous ileal resection (ICR) and the absence of ileal involvement.

**Table 1.** Multivariate logistic regression of low abundance of DP8α and clinical variables in inflammatory bowel disease (IBD) patients. Abbreviations: OR: odds ratio; AZA/6-MP/MTX: Azathioprine/6-mercaptopurine/methotrexate; 5-ASA: 5-aminosalicylic acid; ICR: ileocaecal resection; CRP: C-reactive protein; N/A: not applicable

| Variable | OR | 2.5 % | 97.5 % | P value |
| --- | --- | --- | --- | --- |
| Remission | 0.0346 | 0.0014 | 0.3549 | 0.014 |
| Ileum involvement | 0.0563 | 0.0022 | 0.6556 | 0.041 |
| Gender (male) | 0.1378 | 0.0091 | 1.1269 | 0.099 |
| Age | 1.1454 | 1.0441 | 1.3082 | 0.014 |
| Treatments |  |  |  |  |
| Corticosteroid | 0 | N/A | 5.0720E+087 | 0.991 |
| AZA/6-MP/MTX | 0.1276 | 0.0112 | 0.8611 | 0.054 |
| 5-ASA | 246676327952.358 | 0 | N/A | 0.992 |
| Vedolizumab | 0.0009 | 0 | 0.0685 | 0.009 |
| Ustekinumab | 0 | N/A | N/A | 0.998 |
| ICR | 97.0197 | 2.4363 | 23611.03 | 0.042 |
| Smoker | 30.7739 | 0.9592 | 3238.4917 | 0.081 |
| CRP > 5 | 13.7857 | 1.3577 | 236.7969 | 0.041 |

As previously demonstrated (22), we confirmed that the frequency of CCR6^+^/CXCR6^+^ DP8α per 10,000 CD3^+^ T cells was decreased in CD and UC patients, as compared to healthy controls in the whole cohort (Figure 4A) and also in the subcohort of patients with available fecal microbiota data (Figure 4B). This decrease was seen in both UC and CD, independently of ileal involvement. The quantity of fecal *F. prausnitzii* was also significantly decreased in all groups of patients with IBD, as compared to controls, although the signal was stronger in patients with CD and ileal involvement (Figure 4C), consistent with previous findings (30). Interestingly, we observed that the number of circulating CCR6^+^/CXCR6^+^ DP8α, previously described as specific for *F. prausnitzii*, was positively correlated with the abundance of *F. prausnitzii* in the fecal microbiota of healthy controls and CD patients with ileal involvement but not in purely colonic CD and UC patients (Figure 4D-F).

**Figure 4.**
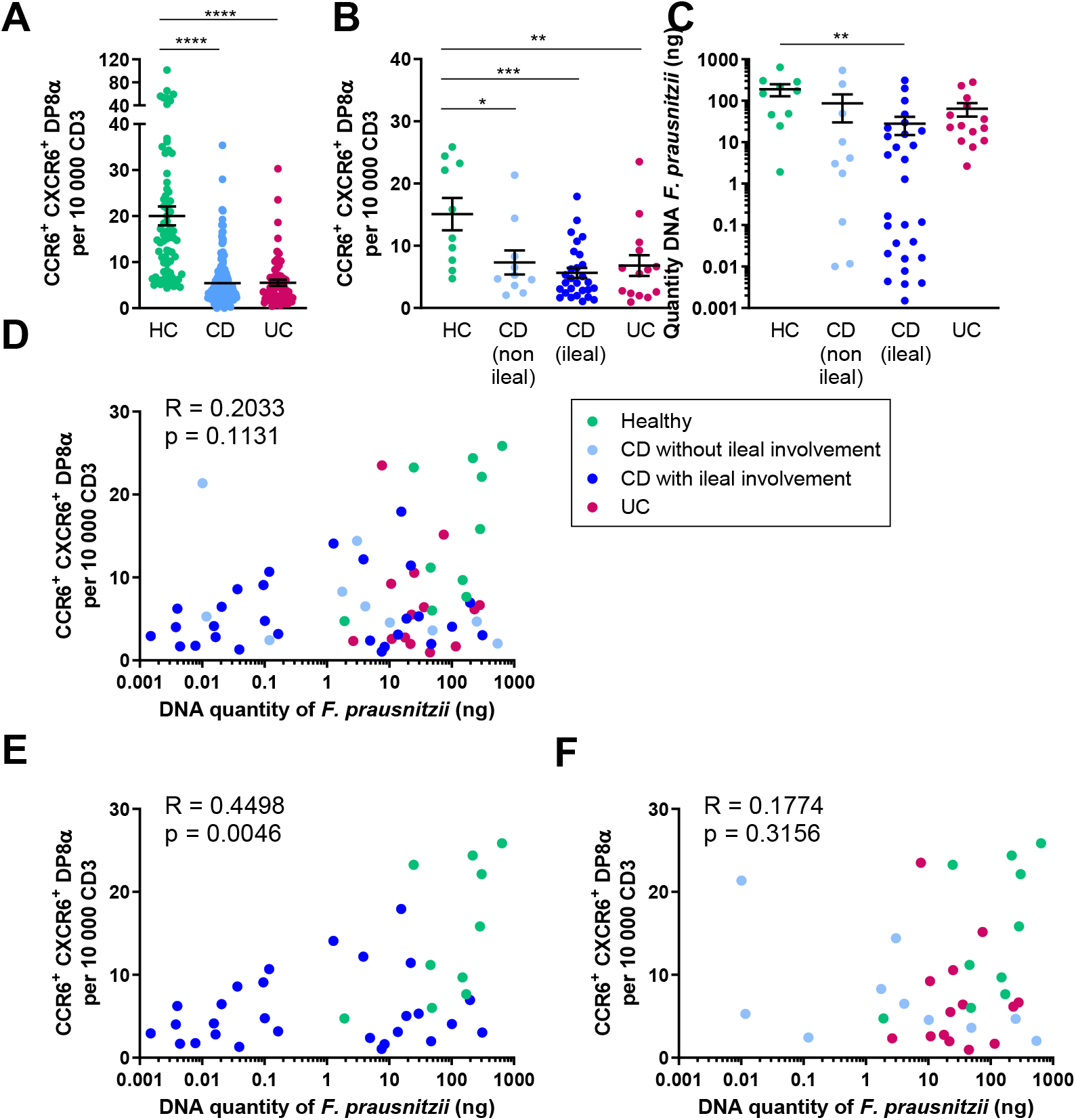
The frequency of circulating CCR6^+^/CXCR6^+^ DP8α Tregs and the quantity of *F. prausnitzii* DNA in fecal microbiota are both decreased in IBD patients and are positively correlated in Crohn’s Disease patients with ileal involvement. **(A)** Frequency of circulating CCR6^+^/CXCR6^+^ DP8α per 10,000 CD3 T cells in the whole cohort of healthy subjects (HC, n = 73), Crohn’s disease patients (CD, n = 185) and Ulcerative Colitis (UC, n = 65) patients, or **(B)** according to localization (with or without ileal involvement) in a subcohort of HC (n = 10) and IBD patients (n = 10 CD without ileal involvement, n = 28 CD with ileal involvement and n = 14 UC patients). **(C)** Quantity of *F. prausnitzii* DNA in stool samples in HC and IBD patients. **(D, E, F)** Spearman correlation of the frequency of circulating CCR6^+^/CXCR6^+^ DP8α cells per 10,000 CD3 T cells and the quantity of *F. prausnitzii* DNA in fecal microbiota in the different groups. Results in panels A-C are presented as the mean ± S.E.M, **P < 0.01, ***P < 0.001, ****P < 0.0001 one-way analysis of variance (ANOVA) followed by Tukey’s multiple comparison tests.

## DISCUSSION

In this study, we demonstrated the protective role of DP8α Tregs *in vivo* in the development of DSS-induced colitis in a sophisticated model of immunodeficient humanized mice. In presence of CD4^+^ effector T cells, the administration of DR*0401-restricted DP8α Treg cells in combination with *F. prausnitzii* intragastric gavage attenuated colitis severity as assessed by the decrease of the disease activity index, body weight loss, lipocalin-2 concentration in colon content and RNA expression of the antimicrobial molecules *Reg3b* and *Reg3g* in the colon of NSGAb°DR*0401 mice, while the administration of DP8α Tregs or *F. prausnitzii* alone had no protective effect. This underlines the importance of *F. prausnitzii*-stimulation to activate DP8α Treg cells to exert their anti-inflammatory role and suggest that these Tregs could play a role in intestinal homeostasis and in the control of intestinal inflammation. We had previously demonstrated the protective effect of *F. prausnitzii* and its supernatant in a model of Trinitrobenzene sulfonic acid (TNBS)-induced acute colitis model (Sokol et al., 2008) and in a model of chronic colitis using Dinitrobenzene sulfonic acid (DNBS) (Martín et al., 2014). However, it is difficult to compare those studies to the present one, as the colitis model and mouse strains are different. In other models of colitis, the effect of *F. prausnitzii* was not clearly demonstrated. Kawade *et al*. showed that live but not inactivated *F. prausnitzii* was able to prevent the effect of DSS-induced colitis in a model of BALB/c mice (Kawade et al., 2019), and another study showed that it is the administration of *F. prausnitzii* supernatant that ameliorates DSS colitis in C57BL/6J mice and that this effect is mediated by Th17 cells. In the present study, we used *F. prausnitzii* in particularly immunodeficient mice with defects in innate immunity and lacking adaptive immune cells, in which the only adaptive immune cells present are therefore the human CD4 effector T cells and DP8α Tregs administered intraperitoneally. So, the system we set up is designed to specifically test the effect of DP8α Tregs. In humans, it is likely that *F. prausnitzii* acts through different mechanisms and targets, including DP8α cells but also epithelial cells (Lenoir et al., 2020; Quévrain et al., 2016) and antigen presenting cells likely responsible for DP8α Treg induction/priming (Alameddine et al., 2019).

To extend our knowledge regarding the clinical relevance of DP8α abundance and its link with *F. prausnitzii,* we took advantage of a large cohort of 250 IBD patients. Interestingly, we observed that a low frequency of circulating total DP8α cells was associated with several parameters of disease activity (including flare status and elevated CRP) but also with previous ileocecal resection and the absence of ileal involvement. These findings suggest that CD patients with severe terminal ileum involvement who required ileocecal resection and thus displayed a more severe phenotype, have fewer DP8α cells. Although we observed that *F. prausnitzii* abundance was decreased in all patients with IBD, patients with CD and ileal involvement exhibited the lowest levels of *F. prausnitzii*, consistent with previous findings (Willing et al., 2009). We also observed a high heterogeneity in the quantity of *F. prausnitzii* in the group of patients with ileal involvement, with two distinct groups of patients (high and low *F. prausnitzii*) that is not entirely explained by localization of the disease (ileal or ileocolonic), previous surgery or activity of the disease. In 2008, we showed that a low proportion of *F. prausnitzii* on resected ileal Crohn mucosa constituted a risk factor for endoscopic recurrence at 6 months (Sokol et al. 2008). Low abundance of *F. prausnitzii* has been confirmed in numerous CD populations ever since, but some studies indicate that this mostly applies to patients with ileal CD (Willing et al., 2009, Willing et al., 2010).

To evaluate the potential link between *F. prausnitzii* and the induction of DP8α cells in humans, we analyzed the correlation between the number of *F. prausnitzii*-specific DP8α T cells, identified by the surface expression of CCR6 and CXCR6 (Godefroy et al., 2018), and the amount of fecal *F. prausnitzii*. We observed a statistically significant positive correlation between both parameters only in healthy individuals and CD patients with ileal involvement but not in purely colonic CD or UC patients. This might be due to the strongest decrease in *F. prausnitzii* in CD patients with ileal involvement that allow to detect more distinctly correlations with DP8α cells abundance.

A main limitation of the present study relies in the rather low strength of the association between circulating DP8α Tregs and fecal *F. prausnitzii* (r value 0.45). The relationship between both variables is probably highly complex and depends on multiple different factors. This correlation might be strengthened by increasing the number of patients, unfortunately, we were able to collect stool samples from only 20.8% of the initial cohort of 250 IBD patients. It would also be of major interest to study the association between ileocolonic lamina propria DP8α and *F. prausnitzii* in IBD patients, since circulating DP8α cells do not necessarily represent their intestinal counterparts. However, our data support the protective role of DP8α Tregs observed *in vivo* in DSS-induced colitis and highlight the potential role of *F. prausnitzii* as a direct inducer of mucosal DP8α cells in humans.

In conclusion, we showed that DP8α cells activated by *F. prausnitzii* protect against intestinal inflammation *in vivo*. The detailed underlying mechanisms remain to be deciphered. The association between low circulating levels of DP8α cells, disease activity and inflammation support the anti-inflammatory role of these cells in humans. Although it does not prove any causal relationship, the association between blood *F. prausnitzii*-specific CCR6+/CXCR6+ DP8α cells and the abundance of fecal *F. prausnitzii* in ileal CD, support the role of *F. prausnitzii* in inducing DP8α in humans with a potential role in gut homeostasis. Supplementing the gut microbiota of IBD patients with *F. prausnitzii,* notably to induce DP8α could therefore represent a promising therapeutic strategy.

## METHODS

### Human subjects

Peripheral blood and stool samples were obtained from healthy individuals and individuals with IBD which were prospectively recruited between 2018 and 2020 in the Gastroenterology Department of the Saint Antoine Hospital (AP-HP, Paris, France). Clinical characteristics and treatments of patients recruited to the study can be found in **Supplemental Table 1 and Supplemental Table 2**. Human Peripheral blood mononuclear cells (PBMCs) derived from healthy volunteers who gave informed consent, at the Etablissement Français du Sang (EFS, Pays de Loire, France) were isolated by Ficoll gradient centrifugation, and used to purify and clone CD4^+^ T cells and DP8α Treg clones, respectively, as detailed in details below and were also used as healthy controls for IBD patients.

### Culture of Faecalibacterium prausnitzii A2-165

*Faecalibacterium prausnitzii* strain A2-165 (DSMZ collection, Braunschweig, Germany) (DSM N° 17677) was grown overnight at 37°C in anaerobic conditions (90% N_2_, 5% CO_2_ and 5% H_2_) in LyBHI [brain-heart infusion medium supplemented with 0.5% yeast extract (Difco Laboratories) and 5 mg/liter hemin] supplemented with cellobiose (1 mg/ml; Sigma Aldrich), maltose (1 mg/ml, Sigma Aldrich), and cysteine (0.5 mg/ml, Sigma Aldrich). A2-165 was pelleted, resuspended in PBS and stored at −80°C. A2-165 was oxygen-inactivated before administration to mice.

### Mouse model

NOD.Cg-Prkdc^scid^ Il2rg^tm1Wj1^ H2-Ab1^tm1^Gru Tg(HLA-DRB1)31Dmz/SzJ (NSG-Ab°DR4) are immunodeficient mice that lack murine T cells, B cells and NK cells, humanized for the expression of a human MHC class II allele (HLA-DR4) in absence of murine MHC II. The animals were purchased from The Jackson Laboratories and housed and bred under Specific and Opportunistic Pathogen Free conditions at the Saint-Antoine Research Center (Plateforme d’Hébergement et d’expérimentation animale, PHEA; Centre de Recherche Saint-Antoine, CRSA; INSERM UMR_S938) and fed *ad libidum.* 10- to 12-week-old female mice were assigned to the experimental groups after matching age and weight and the animals were group housed in the same cage.

### Human cell isolation, cloning and expansion

Human Peripheral blood mononuclear cells (PBMCs) were isolated by Ficoll gradient centrifugation from healthy donor blood (Etablissement Français du Sang, EFS, Pays de Loire, France) and HLA typing was performed (EFS, Pays de Loire, France). CD4 T cells from HLA-DRb1*04-positive donors were magnetically purified using CD4 microbeads according to the supplier’s instructions (Miltenyi Biotec), while CD3^+^/CD4^+^/CD8α^low^/CCR6^+^/CXCR6^+^ *F. prausnitzii-specific* DP8α Tregs were sorted and cloned using a FACS Aria III. CD4^+^ T cells and DP8α Treg clones were expanded on feeder cells, as we previously described (Godefroy et al., 2018).

### DSS-induced colitis in humanized mice

The 10-12 weeks old NSG-Ab°DR4 mice were injected intraperitoneally with 2 x 10^6^ human peripheral CD4 effector T cells isolated from a healthy donor that was positive for the HLA DR4 allele alone or in combination with 2 x 10^6^ DP8α clone DR4/DR4. Following the injection of human cells, the mice received daily intragastric gavages with either 200 μl of PBS alone or 200 μl of A2-165 in PBS (approximately 1 x 10^8^ CFU/mouse/day) for 10 days, before the start of colitis. To induce colitis, mice were administered dextran sulfate sodium (DSS 36,000 - 50,000 Da; MP Biomedicals) at 1-1.5% in drinking water *ad libidum* for 7 days and were then sacrificed or allowed to recover by drinking unsupplemented water for 4 days. In all experiments, body weight, blood in the stool, and stool consistency were analyzed daily. The severity of colitis was assessed using the disease activity index (DAI) with the modified method of Cooper and colleagues (32). Diarrhea was scored as follows: 0, normal; 2, loose stools; 4, watery diarrhea. Blood in stool was scored as follows: 0, normal; 2, slight bleeding; 4, gross bleeding. Weight loss was scored as follows: 0, none; 1–15%; 2, 5–10%; 3, 10–15%; 4, <15%. DAI was the average of these scores.

### Whole blood staining, isolation of LP cells and flow cytometry staining in mice

Whole blood from NSG-Ab°DR4 mice was collected from the submandibular vein before the start of DSS treatment. 100 μl of whole blood was stained with APC labeled anti-mouse CD45, APC-Vio770 labeled anti-human CD45, PE-Vio770 labeled anti-human CD3 and Vioblue labeled anti-human CD4 (all from Miltenyi Biotec, Bergisch Gladbach, Germany) for 10 minutes at room temperature, then incubated with red blood cell lysis solution (Miltenyi Biotec) for 20 minutes and fixed in 1% paraformaldehyde solution for 20 minutes. Finally, the cells were washed and resuspended in PBS 0.5% BSA, 2mM EDTA before acquisition. For isolation of lamina propria (LP) immune cells, the colon was washed with cold PBS and minced into 0.5cm long pieces before incubation in chelating buffer Hanks’ Balanced Salt Solution (HBSS) (Sigma Aldrich) with 5% foetal bovine serum (FBS, PAA Laboratories), 5mM EDTA (Sigma Aldrich), 1mM 1,4-Dithiothreitol (DTT, Sigma Aldrich) for 20 min at 37°C under agitation. The lamina propria was cleaned of epithelial cells after washes with PBS and filtration through 100 μm mesh cell strainer. Cells of the LP were isolated from fibrous matter using collagenase IV (680 UI/ml, Roche) and DNAse I (1 mg/ml, Sigma Aldrich) digestion for 30 min at 37°C under agitation. Cells were washed in Roswell Park Memorial Institute medium (RPMI) 1640 medium (Gibco) supplemented with 10% FBS, 1% penicillin streptomycin (Gibco), 10mM Hepes (Gibco) and 50μM betamercaptoethanol (Sigma Aldrich). LP immune cells were enriched at the interface of a 40%/80% Percoll gradient (GE Healthcare Life Sciences), washed and stained for flow cytometry with the same antibodies used in whole blood, with addition of eFluor 506 fixable viability dye (Thermo Fisher) to assess cell death. Data were acquired on a CytoFLEX (Beckman Coulter) flow cytometer and data were analyzed on FlowJo V10 (Tree Star).

### qRT-PCR in murine colon

Proximal colon tissues from NSG-Ab°DR4 mice were harvested after 7 days of DSS treatment and freezed in liquid nitrogen. Tissues were transferred in Lysing Matrix D tubes (MP Biomedicals) and homogenized on a FastPrep (MP Biomedicals) bead beating machine. RNA was extracted from homogenized tissues using RNeasy Mini Kit (Qiagen) according to the manufacturer’s instructions. RNA samples were reversed-transcribed using High-Capacity cDNA Reverse Transcription Kit (Thermo Fisher). qPCR were performed using SYBR™ Green PCR Master Mix (Applied Biosystems) in a StepOnePlus apparatus (Applied Biosystems). The oligonucleotides used were as follows: *Rplp0 —* sense: 5’-AGATTCGGGATATGCTGTTGGC-3’; antisense: 5’-TCGGGTCCTAGACCAGTGTTC-3’, *Reg3b* — sense: 5’-ATGCTGCTCTCCTGCCTGATG-3’; antisense: 5’-CTAATGCGTGCGGAGGGTATATTC-3’, *Reg3g* — sense: 5’-TTCCTGTCCTCCATGATCAAAA-3’; antisense: 5’-CATCCACCTCTGTTGGGTTCA-3’. The 2-ΔΔCt quantification method was used for the analyses with mouse *Rplp0* as an endogenous control and normalization to CD4 + PBS-treated mice.

### Quantification of colon lipocalin (Lcn-2) levels

During sacrifice and sample collection, colon contents were collected from NSG-Ab°DR4 mice and frozen for further analyses. Frozen colon contents were weighted and suspended in PBS. Samples were vortexed for 20 min to get a homogenous suspension and centrifuged for 10 min at 10,000g at 4°C. Clear supernatants were collected ad stored at −80°C until analysis. Lcn-2 levels were measured using Duoset murine Lcn-2 ELISA Kit (R&D systems) according to the manufacturer’s instructions and results were expressed as pg/g of colon content.

### Histological score

Colon fragments taken from mid-part of the colon were fixed in 4% paraformaldehyde solution, embedded in paraffin and 5mm sections were stained with hematoxylin and eosin for histological scoring. Tissues were scored blindly using established methods for DSS colitis as previously described (4). Inflammation severity, inflammation extent and crypt damage were graded. For each feature, the product of the grade and the percentage involvement was established. The histological score was obtained by adding the subscore of each feature.

### Flow cytometry to track circulating DP8α Tregs in patients and healthy controls

Human peripheral blood mononuclear cells (PBMCs) were isolated by Ficoll gradient centrifugation. After isolation, PBMCs were stained for 45 minutes at 4°C in PBS 0.1% bovine serum albumin with the following antibodies: anti-human CD3-PE-Cy7 (clone UCHT1, Becton Dickinson), anti-human CD4-FITC (clone 13B8.2, Beckman Coulter), anti-human CD8a-BV421 (clone RPA-T8, Becton Dickinson), anti-human CCR6-PE (clone G034E3, Biolegend) and anti-human CXCR6-APC (clone K041E5, Biolegend). Fluorescence was measured on a BD LSR II flow cytometer (BD Biosciences) and analyzed using FlowJo or DIVA softwares.

### DNA extraction and bacterial quantification in fecal microbiota of patients and healthy controls

Fecal genomic DNA from patients was extracted from the stool samples using a method that was previously described (4), which is based on the Godon DNA extraction method (33). More precisely, the feces samples were weighed and then resuspended for 10 min at room temperature in 250 μl of 4 M guanidine thiocyanate in 0.1 M Tris-HCl (pH 7.5) (Sigma-Aldrich) and 40 μl of 10% N-lauroyl sarcosine (Sigma-Aldrich). After the addition of 500 μl of 5% N-lauroyl sarcosine in 0.1 M phosphate buffer (pH 8.0), the 2-ml tubes were incubated at 70 °C for 1 h. One volume (750 ml) of 0.1-mm-diameter silica beads (Sigma-Aldrich) (previously sterilized by autoclaving) was added, and the tube was shaken at 6.5 m/s three times for 30 s each in a FastPrep (MP Biomedicals) apparatus. 15 mg of polyvinylpolypyrrolidone (Sigma) was added to the tube, which was then vortexed and centrifuged for 5 min at 20,000g at 4°C. After recovery of the supernatant, the pellets were washed with 500 μl of TENP (50 mM Tris (pH 8), 20 mM EDTA (pH 8), 100 mM NaCl, 1% polyvinylpolypyrrolidone) and centrifuged for 5 min at 20,000g at 4°C, and the new supernatant was added to the first supernatant. The washing step was repeated two times. The pooled supernatant (about 2 ml) was briefly centrifuged to remove particles and then split into two 2-ml tubes. Nucleic acids were precipitated by the addition of 1 volume of isopropanol for 10 min at room temperature and centrifugation for 10 min at 20,000g at 4°C. Pellets were resuspended and pooled in 450 μl of 100 mM phosphate buffer, pH 8, and 50 μl of 5 M potassium acetate. The tube was placed at 4°C overnight and centrifuged at 20,000g for 30 min. The supernatant was then transferred to a new tube containing 20 μl of RNase (1 mg/ml) and incubated at 37 °C for 30 min. Nucleic acids were precipitated by the addition of 50 μl of 3 M sodium acetate and 1 ml of absolute ethanol. The tube was incubated for 10 min at room temperature, and the nucleic acids were recovered by centrifugation at 20,000g for 15 min. The DNA pellet was finally washed with 70% ethanol, dried, and resuspended in 100 μl of Tris–EDTA (TE) buffer. DNA suspensions were stored at −20 °C for real-time quantitative polymerase chain reaction (qPCR) analysis. Quantifications of *F. prausnitzii* were performed by qPCR using SYBR™ Green PCR Master Mix (Applied Biosystems) in a StepOnePlus apparatus (Applied Biosystems). Each reaction was done in duplicate in a final volume of 20 μl wih 0.2 μM of each primer and 5 μl of the appropriate dilution of DNA. *F. prausnitzii* was quantified using specific primers: (sense) 5’-CCATGAATTGCCTTCAAAACTGTT-3’ and (antisense) 5’-GAGCCTCAGCGTCAGTTGGT-3’ (13). Amplifications were performed with the following conditions: denaturation step 10 min at 95°C; 40 cycles of 95°C denaturation for 15 sec; 55°C extension for 1 min followed by a standard melting curve program. We used quantified *F. prausnitzii* DNA as reference standard to quantify *F. prausnitzii* DNA in each sample.

### Statistics

GraphPad Prism version 6.0 (GraphPad, La Jolla, CA) was used for statistical analysis in mouse experiments, comparisons of DP8α frequencies or abundance of *F. prausnitzii* between healthy donors and patients and for all graphs representations. For data displayed in graphs, the results are presented as the mean ± S.E.M. For comparison between multiple groups, one-way analysis of variance (ANOVA) was performed, and P value < 0.05 were considered significant (*P <0.05, **P < 0.01, ***P < 0.001, ****P < 0.0001 and § P<0.05, §§ P < 0.01). Only significant statistical results after adjustment for False discovery rate are shown (Benjamini Hochberg, q< 0.1). Spearman correlations were used for correlations between DP8α frequencies and abundance of fecal *F. prausnitzii* in healthy controls and patients. Statistical analysis on the patients’ cohort was performed in the R statistical environment (R version 3.6.2). For multivariate logistic regression with stepwise variable selection, the variables taken into consideration were: gender, age, disease (UC/CD), activity (flare/remission), topography (ileum, colon), treatment, history of surgery (ileocecal resection, colectomy, ileostomy), smoking (active or not), CRP (elevated or normal).

### Study approval

All human samples were collected with informed consent. Approval for human studies was obtained from the local ethics committee (Comité de Protection de Personnes Ile de France IV, IRB 00003835 Suivitheque study; registration number 2012/05NICB). For isolation of PBMC to purify and clone CD4^+^ T cells and DP8α Treg clones, blood samples were obtained from the Etablissement Français du Sang (EFS, Pays de Loire, France) (Convention number CPDL-PLER-2017 21). All animal experiments were carried out in strict accordance with the French national and European laws and conformed to the Council Directive on the approximation of laws, regulations, and administrative provisions of the Member States regarding the protection of animals used for experimental and other scientific purposes (86/609/Eec). All animal experiments were conducted according to the institutional guidelines approved by the local ethics committee (reference number APAFIS#8231-2016121612402727 v2).

## Supporting information

supplemental informations and figures

## AUTHOR CONTRIBUTIONS

Conceptualization, FJ, FA and HS.; Methodology, ST and EG.; Investigation, ST, EG, NR, CD, CO, MS, CG, LB, IAS, TL and SC; Formal Analysis, ST and HS.; Writing – Original Draft, ST, NR and HS; Writing – Review & Editing, ST, EG, NR, FJ, and FA and HS; Funding Acquisition, FA, FJ and HS; Resources, TL (mice), JMC and PL (bacteria), HS (human data and samples).; Supervision, FJ, FA and HS.

## ACKNOWLEDGMENTS

The authors thank the patients recruited in the Gastroenterology Department of Saint-Antoine Hospital (AP-HP, Paris, France) and the nurses, the members of the animal core facility (PHEA, Saint-Antoine Research Center), B. Solhonne (Saint-Antoine Research Center, Paris, France) for her contribution for histology staining, Pr M. Mohty and Dr B. Gaugler (Saint-Antoine Research Center, Paris, France) for the use of the CytoFLEX flow cytometer and M-L Michel (INRAE, Jouy-en-Josas, France) for precious advice on murine lamina propria isolation.

