## supplemental informations and figures for "Human CD4^+^/CD8α^+^ regulatory T cells induced by *Faecalibacterium prausnitzii* protect against intestinal inflammation"

**SUPPLEMENTAL INFORMATION**

**Supplemental Methods**

*Screening for HLA-restriction.* Grown DP8α Treg clones were screened for their HLA-DR restriction using the L243 blocking antibody. To this end, autologous monocytes ($CD14^+$ magnetically purified, Miltenyi Biotec) were loaded overnight with *F. prausnitzii* (ratio 1 monocyte:1 bacterium) before addition of individual DP8α Treg clones (ratio 2 clones:1 monocyte), in the presence or in the absence of the L243 antibody (10µg/ml) or its corresponding control Ig. Selected HLA-DR*04-restricted clones were then extensively expanded for *in vivo* experiments.

*Isolation of murine bone-marrow derived dendritic cells.* For co-culture with DP8α Treg clones, bone-marrow derived dendritic cells from a 10 week-old NSG-Ab°DR4 mice were differentiated *ex vivo*. Femurs and tibias were collected from the mouse, soaked in 70% ethanol for 5 min and placed in Iscove's Modified Dulbecco's Medium (IMDM) medium supplemented with 10% FBS, 1% L-Glutamine (Gibco), 1% Penicillin/streptomycin (Gibco), 50 µM of betamercaptoethanol (Sigma) (complete medium). Bone marrow was flushed from the bones using a 2 ml syringe and 26G needle, collected in a 50 ml tube and centrifugated 5 minutes at 300g at 4. Supernatant was discarded and the cells were incubated at room temperature with red blood cell lysis solution (Miltenyi Biotec) for 5 minutes. After centrifugation 5 minutes at 300 g at room temperature, the cells were washed and resuspended in complete medium supplemented with 10 ng/ml recombinant mouse granulocyte-macrophage colony-stimulating factor (GM-CSF; Miltenyi Biotec) at the concentration of 5 x $10^6$ cells / ml in non-culture treated bacterial petri dish (Sarstedt) and placed at 37°C 5% $CO_2$ for 5 days. At day 5, culture medium was replaced. At day 7, BM-DC were detached using cold PBS, counted and frozen in FBS + 10% DMSO before being used for co-culture experiments.

*Human IL-10 and IFNɣ ELISAs.* DP8α Treg clones were stimulated either specifically by APCs: autologous monocytes or murine NSG-Ab°DR4 BM-DC (ratio 2 lymphocytes:1 APC) loaded overnight or not with the A2-165 *F. prausnitzii* strain (multiplicity of infection, MOI 10), or non-specifically using coated anti-CD3 (clone OKT3, 1 µg/ml, eBioscience, San Diego, CA) for 48h at 37°C. Supernatants were harvested at 24h for IFNɣ and 48h for IL-10. Cytokine production was measured using Ready-Set-Go ELISAs (eBioscience) according to the manufacturer’s instructions.

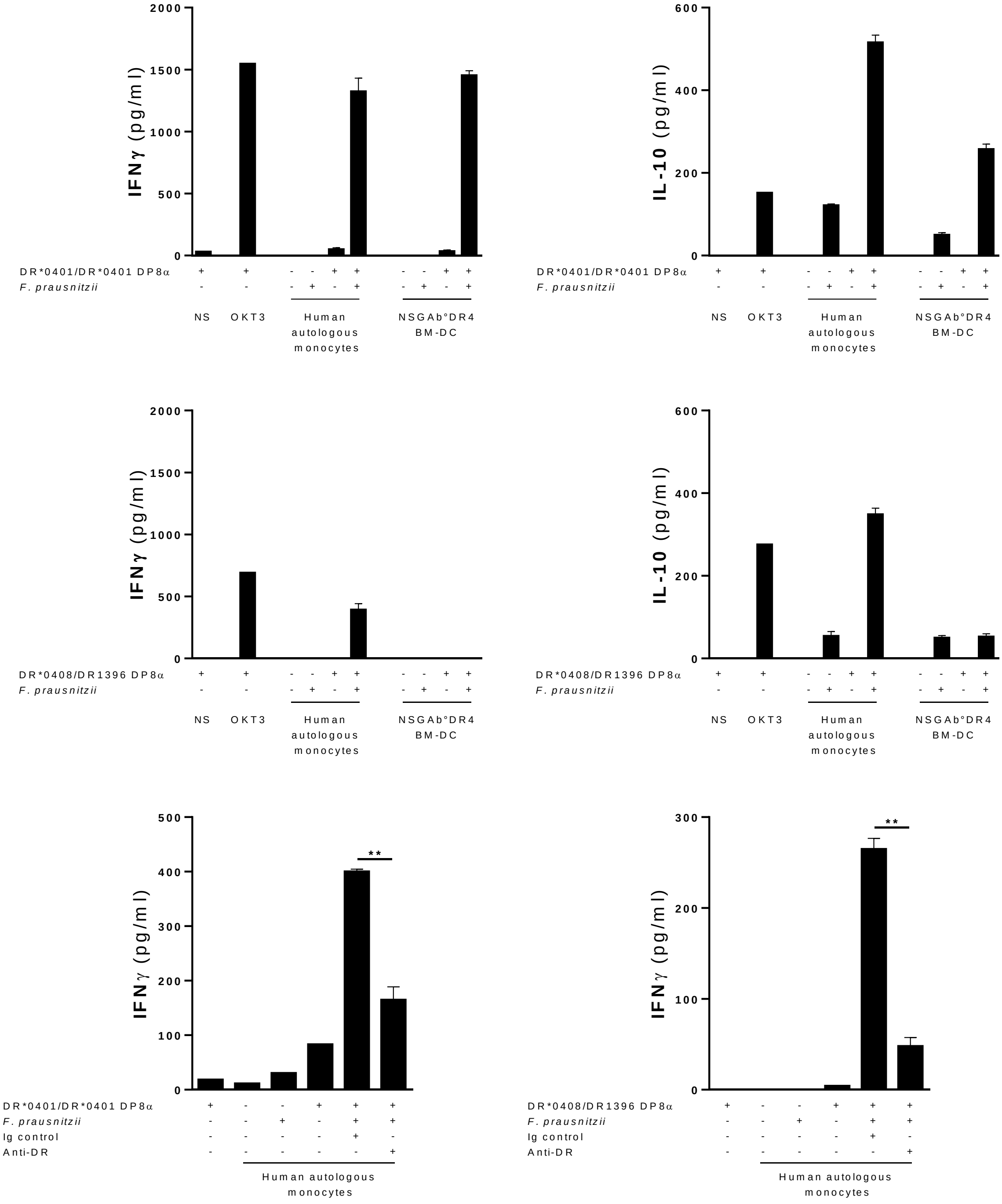


**Supplemental Figure 1. A human DP8α Treg clone from a homozygous DRb1*0401/DRb1*0401 donor responds to bone-marrow derived dendritic cells (BM-DCs) from NSG-Ab°DR4 mice loaded with *F. prausnitzii*. (A, B)** HLA-DR restriction of the DRb1*0401 / DRb1*0401 and the DRb1*0408 / DRb1*1396 DP8α clones using an anti-human HLA-DR blocking antibody (L243 clone) or a control Ig. **(C, D)** IFNɣ and **(E, F)** IL-10 production upon co-culture of DP8α Treg clones in non-stimulating conditions (NS), in response to non-specific stimulation (anti-OKT3 antibody), human autologous monocytes or murine NSG-Ab°DR4 bone marrow derived dendritic cells (BM-DC) loaded or not with *F. prausnitzii*. Results are presented as the mean ± S.E.M (n = 3). Two-tailed Mann-Whitney tests.

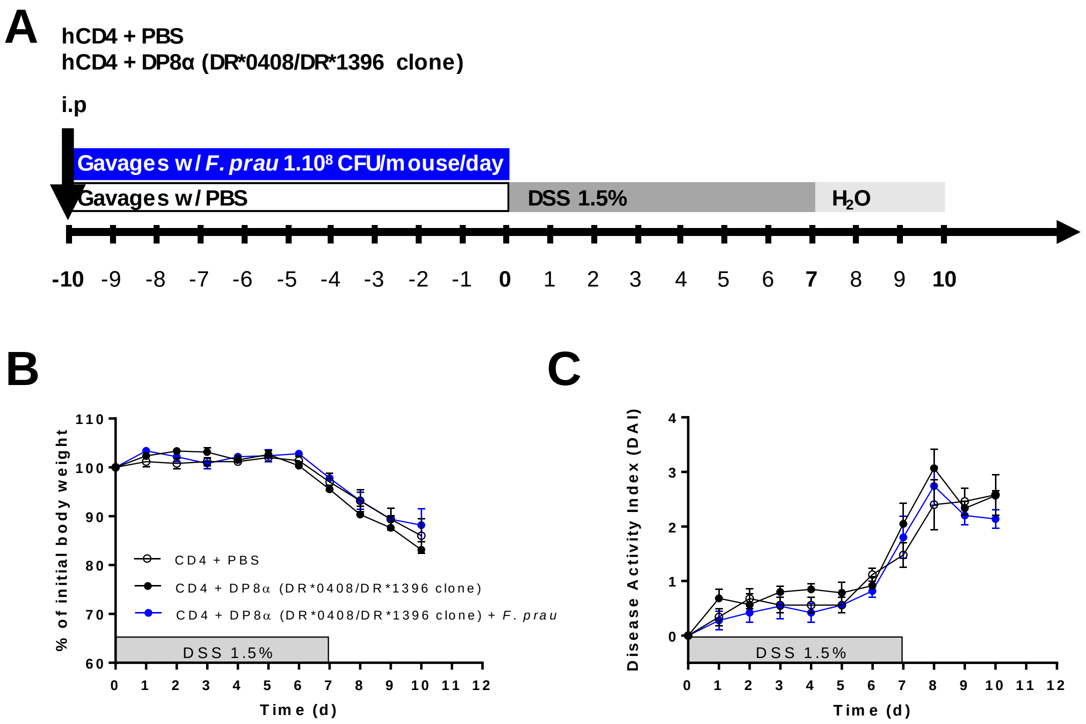


**Supplemental Figure 2. Human DP8α Treg clone from a DRb1*0408 / DRb1*1396 donor has no protective effect on DSS-induced colitis in NSG-Ab°DR4 mice. (A)** Experimental outline: NSG-Ab°DR4 female mice were injected intraperitoneally (i.p.) with $2.10^6$ human peripheral CD4 effector T cells alone, or in combination with $2.10^6$ human DR*0408/DR*1396 DP8α clones and received daily intragastric gavage with 200 µl PBS 1X or $1.10^8$ CFU of *F. prausnitzii* for 10 days before 1.5% DSS supplementation in drinking water for 7 days followed by 3 days of regular drinking water. **(B)** Body weight and **(C)** disease activity index (DAI) were assessed during the protocol in all groups of mice.

| Variable | HC | CD | UC |
|---|---|---|---|
| Group [n] | 73 | 185 | 65 |
| Age [years] | 39 (24.5) | 39 (20) | 36 (17.5) |
| BMI [kg/m$^2$] | - | 22.9 (4.8) | 21.9 (5.4) |
| Gender [male] | 41 (56.2%) | 115 (62.2%) | 37 (56.7%) |
| Intestinal resection | - | 72 (38.9%) | 2 (3.1%) |
| Smoking | - | 40 (21.6%) | 3 (4.6%) |

**Supplemental Table 1. Clinical characteristics of healthy individuals and inflammatory bowel disease (IBD) patients recruited in the study.** Continuous variables are presented as median (interquartile range). Categorical variables are presented as counts (%). Abbreviation: HC: healthy controls, CD: Crohn’s disease, UC: ulcerative colitis, BMI: Body mass index.

| Variable | CD | UC |
|---|---|---|
| Montreal (CD) | | |
| L1 | 35 (18.9%) | - |
| L2 | 42 (22.7%) | - |
| L3 | 72 (38.9%) | - |
| L4 | 1 (0.5%) | - |
| L1 + L4 | 13 (7.0%) | - |
| L3 + L4 | 20 (10.8%) | - |
| Montreal (UC) | | |
| E1 | - | 2 (3.1%) |
| E2 | - | 25 (38.5%) |
| E3 | - | 38 (58.5%) |
| Oral 5'-ASA | 21 (11.35%) | 28 (43.07%) |
| AZA/6-MP | 41 (22.16%) | 18 (27.69%) |
| Methotrexate | 22 (11.89%) | 1 (1.54%) |
| Anti-TNFα | 158 (85.41%) | 53 (81.54%) |
| Vedolizumab | 18 (9.73%) | 12 (18.46%) |
| Ustekinumab | 1 (0.54%) | 0 (0%) |

**Supplemental Table 2. Disease extent, disease severity and medications specific for IBD patients**. Disease extent and medications presented as counts (%). Disease activity presented as median (range). Abbreviation: AZA/6-MP: Azathioprine, 6-mercaptopurine; 5-ASA: 5-aminosalicylic acid
